# Bidirectional Energy Flow in the Photosystem II Supercomplex

**DOI:** 10.1101/2023.11.30.569278

**Authors:** Cristina Leonardo, Shiun-Jr Yang, Kaydren Orcutt, Masakazu Iwai, Eric A. Arsenault, Graham R. Fleming

## Abstract

The water splitting capability of Photosystem II (PSII) of plants and green algae requires the system to balance efficient light harvesting along with effective photoprotection against excitation in excess of photosynthetic capacity ^1,2^, particularly under the naturally fluctuating sunlight intensity. The comparatively flat energy landscape of the multi-component structure, inferred from spectra of the individual pigment-protein complexes and the rather narrow and featureless absorption spectrum, is well known ^3–7^. However, how the combination of the required functions emerge from the interactions among the multiple components of the PSII supercomplex (PSII-SC) cannot be inferred from the individual pigment-protein complexes. In this work, we investigate the energy transfer dynamics of the C_2_S_2_-type PSII-SC with a combined spectroscopic and modeling approach. Specifically, two-dimensional electronic-vibrational (2DEV) spectroscopy ^8,9^ provides enhanced spectral resolution and the ability to map energy evolution in real space, while the quantum dynamical simulation allows complete kinetic modeling of the 210 chromophores. We demonstrate that additional pathways emerge within the supercomplex. In particular, we show that excitation energy can leave the vicinity of the charge separation components, the reaction center (RC), faster than transferring to it. This enables activatable quenching centers in the periphery of the PSII-SC to be effective in removing excessive energy in cases of over-excitation ^2^. Overall, we provide a quantitative description of how the seemingly contradictory functions of PSII-SC arise from the combination of its individual components. This provides a fundamental understanding that will allow further improvement of artificial solar energy devices and bioengineering processes for increasing crop yield ^10^.

## Kinetic network within the complete PSII-SC

In the thylakoid membrane, PSII is bound with light-harvesting complex II (LHCII) trimers to form PSII-LHCII supercomplexes, in a ratio that depends on the acclimated condition. The C_2_S_2_-type PSII-SC (referred to as the PSII-SC in the following) is the dominant form in high-light conditions ^11,12^, where photoprotection is crucial. The arrangement of the chromophores and the protein subunits in the C_2_S_2_-type PSII-SC from spinach is shown in Figure 1a-b. The high-resolution cryo-EM structure ^13^ shows that the PSII-SC is a dimeric complex with 4 pheophytins and 206 chlorophylls (Chls), of which 156 are Chls *a* and 50 are Chls *b*. Each monomer contains one RC, consisting of two branches (D1 and D2), two core antennae (CP43 and CP47), two minor antennae (CP26 and CP29) and one strongly bound LHCII. The subunit containing the RC, CP43 and CP47 is referred to as the PSII core complex (PSII-CC). Among the antennae, LHCII, CP26 and CP43 are in proximity to the D1 branch, the branch that actively performs charge separation ^14,15^, while CP29 and CP47 are on the D2 side of the complex.

**Fig. 1.**
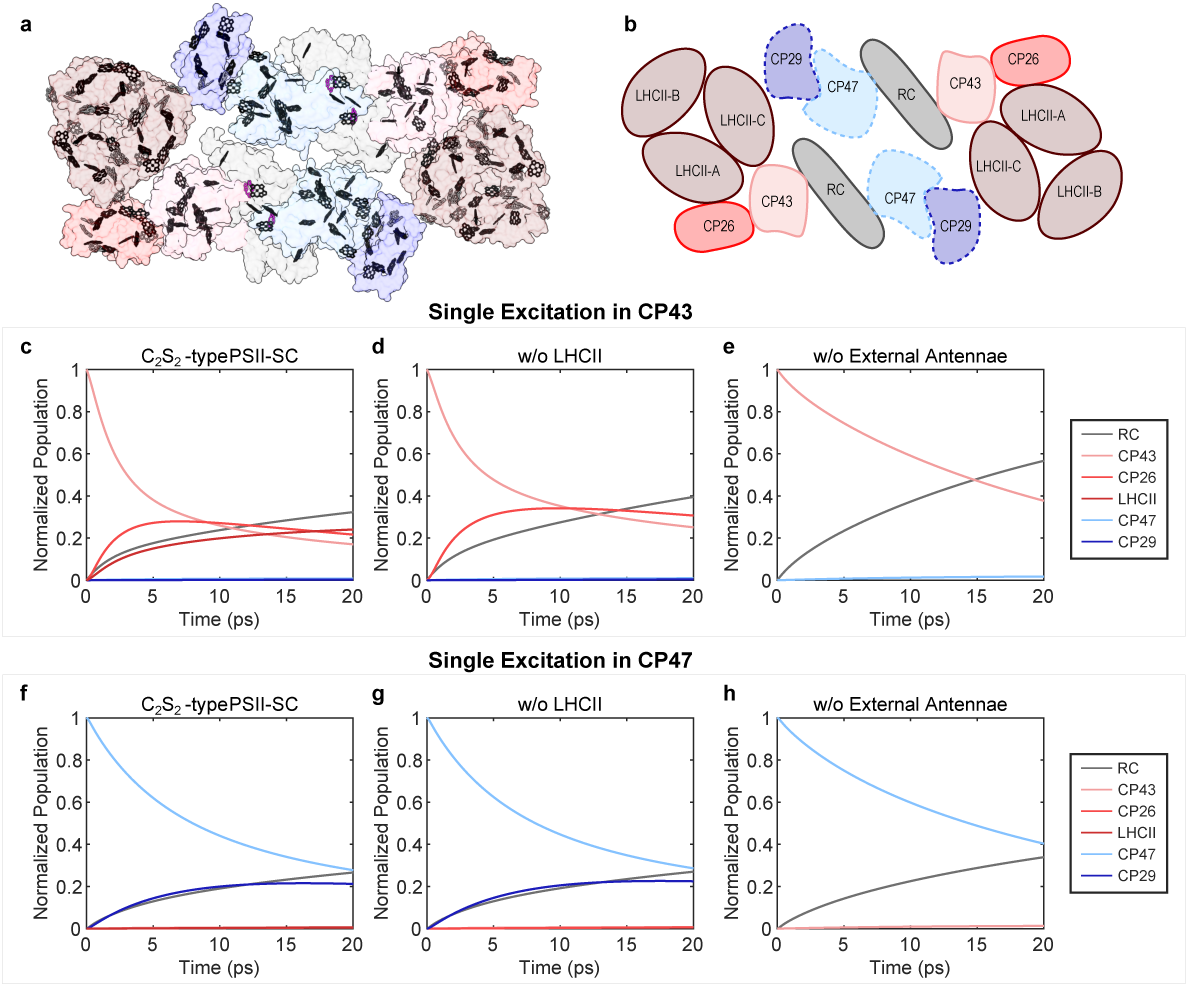
Excitation population traces upon single excitation at 300 K. **a**, Pigment arrangement in the C_2_S_2_-type PSII-SC from spinach (PDB: 3JCU) ^13^: Chls *a* in black; Chls *b* in gray; pheophytins in magenta. **b**, A schematic of the PSII-SC protein subunits. Proteins on the D1 side are represented by solid lines and those on the D2 side are represented by dashed lines. Simulated excitation population dynamics of **c**, the complete C_2_S_2_-type PSII-SC **d**, PSII-SC without LHCII and **e**, without LHCII, CP26 and CP29 upon single excitation in CP43. Simulated excitation population dynamics of **f**, the complete C_2_S_2_-type PSII-SC **g**, PSII-SC without LHCII and **h**, without LHCII, CP26 and CP29 upon single excitation in CP47.

The subunits of the PSII-SC cooperatively form an electronic energy transfer (EET) network that initiates photosynthesis. In particular, the PSII-SC functions strongly rely on the inter-protein EET pathways that originate from its multi-component structure. To understand how the multi-unit construction relates to the photosynthetic functions of PSII, it is necessary to study the complete PSII-SC as some of the crucial pathways are only present when all the subunits are connected. Furthermore, the presence of these pathways alters the significance of the pathways in isolated subunits. We demonstrate this by comparing the EET dynamics obtained from structure-based simulations for the PSII-SC subunits of different sizes. Briefly, we adapt the kinetic model of the PSII-SC proposed by Bennett et al. ^16^ and follow their methods to reconstruct a new structure-based model for the C_2_S_2_-type PSII-SC based on the state-of-the-art high-resolution structure (protein data bank: 3JCU)^13^. The kinetic model was used to produce excitation population evolution, which was coarse-grained to allow focus on the inter-protein EET pathways. Figure 1c-h shows the evolution of the excitation population in the PSII-SC subunits with different sizes of the antenna system upon the excitation of the core, CP43 and CP47. While the initial excitation is placed in the same Chl in either CP43 or CP47, the absence of LHCII and/or the minor antennae alters the EET dynamics. For example, the absence of LHCII results in faster RC growth and slower CP43 decay, while the absence of all external antennae results in even more obvious shifts in timescales. The different traces indicate that the timescales observed in the smaller subunits of the PSII-SC do not necessarily reflect the EET pathways relevant in the native environment, where PSII exists in the form of the PSII-SC. Additionally, potential structural changes caused by the removal of protein subunits, not taken into account in the simulations, can further alter the EET pathways present in the complete system. These stress the importance of studying the entire PSII-SC as the investigation of smaller subunits can lead to inaccurate description of the relevant EET pathways that initiate photosynthesis in nature.

The study of the EET dynamics in the complete PSII-SC is challenging. These dynamics typically occur within tens of fs to a few hundred ps, requiring the use of ultrafast spectroscopic techniques. However, significant spectral congestion due to the large number of Chls present in the PSII-SC challenges the currently available technology. As a result, most studies focus on the smaller subunits of the PSII-SC where the numbers of convoluted processes are reduced ^17–24^. As demonstrated earlier, such studies provide insight into EET pathways existing within the subunits, but are insufficient for obtaining a complete description of energy flow within the PSII-SC relevant to its functions. In the past, fluorescence lifetime studies have been reported for the entire super-complex ^25–29^. However, the energy flow between different subunits cannot be inferred directly from these experiments. Moreover, the uncertainty in kinetic modeling based on the fitting of fluorescence decays cannot be avoided. In addition to the experimental studies, many theoretical simulations have been performed to understand the dynamics within different PSII subunits ^5,7,30–32^. While these works provide detailed understanding of the EET dynamics in each subunit, a full simulation of the PSII-SC is required to connect the microscopic interactions to its ability to balance efficiency and photoprotection. Based on these works, Bennett et al. constructed the structure-based model of the PSII-SC mentioned earlier ^16^. However, the structural information available at the time was not at high resolution (*∼*12 Å) and can only be used to determine relative orientations and approximate distances between individual proteins. Valkunas and coworkers proposed a model that takes into account the heterogeneity of the PSII-SC by introducing excitation diffusion parameters ^33^. The parameters, extracted from fitting fluorescence decays, reveal the connectivity between different protein subunits but do not provide information on specific EET pathways.

To investigate the EET dynamics in the PSII-SC, we rely on a combination of two-dimensional electronic-vibrational (2DEV) spectroscopy^8,9^ and the structure-based dynamical simulation to characterize the early-time (*<*20 ps) inter-protein EET dynamics within the C_2_S_2_-type PSII-SC from spinach. In 2DEV spectroscopy, the simultaneous resolution along the visible excitation and the mid-infrared (IR) probe reduces the spectral congestion that limits the resolution of other ultrafast spectroscopic techniques, thus enabling the study of complex systems such as the PSII-SC ^34–40^. Specifically, the mid-IR detection provides a means to distinguish different protein subunits as the localized vibrational modes are sensitive to the protein surrounding. Meanwhile, the high-resolution cryo-EM structure ^13^ allows a more accurate description of the interactions between the pigments, which leads to a more accurate kinetic model. Together, they reveal the inter-protein EET pathways only present in the entire PSII-SC and are crucial for efficient energy conversion and effective photoprotection.

## Bidirectional energy transfer

Figure 2a shows the 2DEV spectrum of the PSII-SC at a time delay of 120 fs. In the spectrum, positive (red) and negative (blue) features are ground state bleach (GSB) and excited state absorption (ESA) signals, respectively. To avoid photodamage, the experiments were performed at 77 K (see Methods). The observed IR features correspond to the stretching modes of the 13^1^-keto (1,610-1,710 cm^-1^) and the 13^3^-ester carbonyl groups (1,710-1,760 cm^-1^) of Chls *a* and Chls *b* ^18,38,39,41–43^. Both functional groups are good probes of the local protein environment, providing IR markers for the proteins in the PSII-SC. The excitation frequency of 15,200 cm^-1^ marks the separation of two experiments (details in Methods, Extended Data Figure E.1). The lower-frequency excitation range (14,700-15,200 cm^-1^) roughly corresponds to Chl *a* excitation and the higher-frequency excitation range (15,200-15,600 cm^-1^) roughly corresponds to Chl *b* excitation.

**Fig. 2.**
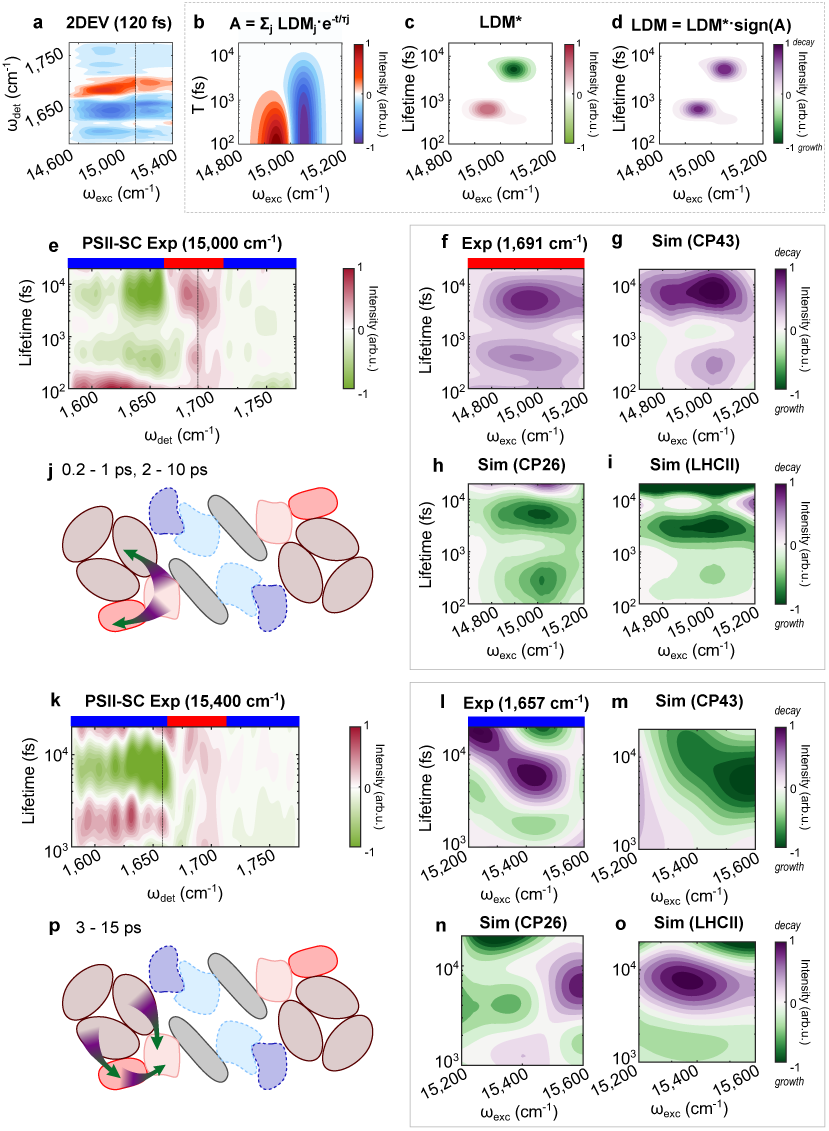
LDM for the PSII-SC at the lower (14,800-15,200 cm^-1^) and the higher excitation frequencies (15,200-15,600 cm^-1^). **a**, The 2DEV spectrum of the PSII-SC at time delays T = 120 fs. Positive (red) and negative (blue) amplitudes are ground state bleach (GSB) and excited state absorption (ESA) signals, respectively. The dashed line marks the separation of two experiments (see Methods). **b**, Simulated evolution of excitation population, where both the GSB and ESA features decay. **c**, Corresponding LDM showing different signs due to the sign difference between GSB and ESA.**d**, LDM multiplied by signal sign, where positive amplitudes indicate decays. **e**, LDM of the 2DEV data averaged over 14,800-15,200 cm^-1^: vertical dashed line at 1,691 cm^-1^; blue (red) bars on top refer to ESA (GSB) signatures. Excitation-dependent LDM of **f**, the GSB at 1,691 cm^-1^ from the 2DEV data and the simulated population evolution of **g**, CP43, **h**, CP26 and **i**, LHCII. Positive amplitudes (purple) indicate decay; negative amplitudes (green) indicate growth. **j**, A schematic of the corresponding inter-protein EET at 0.2-1 ps and 2-10 ps. **k**, LDM from the 2DEV data averaged over 15,300-15,500 cm^-1^: vertical dashed lines at 1,657 cm^-1^; blue (red) bars on top refer to ESA (GSB) signatures. Excitation-dependent LDM of **l**, the ESA at 1,657 cm^-1^ (reversed in sign) and simulated population evolution of **m**, CP43, **n**, CP26 and **o**, LHCII. Positive amplitudes (purple) indicate decay; negative amplitudes (green) indicate growth. **p**, A schematic of the corresponding inter-protein EET at 3-15 ps.

Lifetime density analysis (LDA) is applied to visualize the evolution in the 2DEV spectra and the simulated evolution of the excitation population in each protein (see Methods). LDA provides a way to simultaneously process large parallel data sets, particularly for time-resolved 2D spectra. However, unlike other techniques typically applied for this purpose, such as global and target analysis, LDA does not rely on initial assumptions. Instead, LDA approximates dynamic evolution with hundreds of exponential components to retrieve the amplitude of each lifetime component, leading to bias-free analyses ^44,45^. For different analysis purposes, we use two different types of lifetime density maps (LDMs) to show the pre-exponential factors, i.e., the amplitudes of each exponential component. First, lifetime *vs* detection frequency maps (Figure 2e and k, pink-green maps) are used to identify the principal lifetime components as well as the IR features involved in the main spectral evolution in each excitation frequency range. However, the fact that 2DEV spectra have positive (GSB) and negative (ESA) features complicates the interpretation of LDMs. For example, for ESA features, which have a negative sign, decays have negative amplitudes. This is in contrast to GSB features (positive sign), for which decays have positive amplitudes (Figure 2b-d). Therefore, for GSB features, positive and negative LDM amplitudes indicate decay and growth, respectively. In contrast, for ESA features, positive and negative LDM amplitudes indicate growth and decay, respectively. For this reason, the LDM in Figure 2l for the ESA feature at 1,657 cm^-1^ is shown with a reverse sign so that, in Figure 2f-i and l-o, *all* positive LDM amplitudes represent decays and *all* negative LDM amplitudes represent growths, as specified in the colorbars. These are the second type of LDM, lifetime *vs* excitation frequency maps (purple-green maps), which are used to show the evolution of selected IR features reflecting protein-specific dynamics, as well as the simulated evolution of the excitation population in the corresponding protein.

Figure 2e shows the LDM for the IR frequencies 1,580-1,770 cm^-1^ averaged over excitation frequency 14,800-15,200 cm^-1^. Several IR features decay at 0.2-1 ps and 2-10 ps. Among them, the GSB at 1,691 cm^-1^ is a feature that has been observed in vis-pump IR-probe experiments of isolated CP43 ^19^, as well as in a 2DEV experiment on the PSII-CC ^39^. Moreover, this GSB signature is absent in the vis-pump IR-probe experiment of isolated PSII-RC ^38,43^, isolated CP47 ^18^ and in the 2DEV spectrum of isolated LHCII ^35^ in this excitation frequency range. Since the amino acid residues interacting with the Chls in CP26 and CP29 are very similar to those in LHCII ^13^, we expect similar vibrational signatures for these antennae. The GSB at 1,691 cm^-1^ is, therefore, attributed to CP43. The LDM of this GSB signature (Figure 2f) agrees extremely well with the decays observed for the simulated evolution of the excitation population in CP43 (Figure 2g). Interestingly, it is clear that the simulated LDMs of CP26 and LHCII (Figure 2h-i) show growths on the same timescale as the experimental and simulated CP43 decays. This indicates that inter-protein EET occurs from CP43 to CP26 and LHCII in 0.2-1 ps and in 2-10 ps (Figure 2j). It is important to note that, while the simulation only focuses on inter-protein dynamics, the sub-ps decay of CP43 observed experimentally may also have a contribution from intra-protein EET within CP43, as observed in the isolated PSII-CC ^39^. However, the intra-protein EET in CP43 occurs on a slightly shorter timescale (*∼*180 fs) than the observed timescale (0.2-1 ps) which suggests that most contribution comes from inter-protein EET from CP43 to CP26 and LHCII. Overall, the two timescale ranges observed for the CP43 to CP26/LHCII transfer are much shorter than the reported values for the EET from the core antennae to the RC (which range from 10-50 ps in different studies ^20,26,30,39,46^).

Figure 2k shows the experimental LDM averaged over excitation frequency 15,300-15,500 cm^-1^ for the PSII-SC, highlighting two main dynamics: EET around 1-3 ps and 3-15 ps. To show the origin of these dynamics, we select the ESA at 1,657 cm^-1^, the strongest signature in the averaged LDM. The ESA feature at 1,657 cm^-1^ is observed only for the higher-frequency excitation range in the 2DEV data of the PSII-SC, for which mostly Chls *b*—only present in the peripheral antennae—are excited. It is not observed in the 2DEV slices of the PSII-CC (Extended Data Figure E.1). Therefore, this signature is attributed to the peripheral antennae. In fact, the 2DEV slices at 120 fs for the excitation frequencies of 15,300 and 15,600 cm^-1^ for the PSII-SC (Extended Data Figure E.1h, blue and dark blue) have an ESA feature between the detection range 1,630-1,660 cm^-1^ that resembles the ESA feature in the same detection frequency range for isolated LHCII (Extended Data Figure E.1i, blue and dark blue). The ESA feature is also clearly distinct from that of the lower excitation frequencies of the PSII-SC (Extended Data Figure E.1h, dark red and red) and the PSII-CC (Extended Data Figure E.1j, dark red and red), which resemble each other. All the evidence shows that the strong ESA peak at 1,657 cm^-1^ is a marker for the peripheral antennae. In particular, LHCII has a much larger number of Chls compared to CP26 and CP29, and therefore, the contribution from the minor antennae to this signature is expected to be small. The LDM associated with the ESA at 1,657 cm^-1^ is shown in Figure 2l. For comparison, the LDMs of the simulated evolution for CP43, CP26 and LHCII are shown in Figure 2m-o. It is clear that the evolution of the ESA signature shows strong agreement with the simulated evolution of LHCII, further strengthening our assignment of the ESA at 1,657 cm^-1^ to the major antenna. More complex patterns are observed in this excitation frequency range (15,200-15,600 cm^-1^), where mostly Chls *b* (found only in the external antennae) are excited. First, the excitation population in LHCII grows within 1-3 ps, while the only excitation population decay observable within the same timescale is found in CP26, indicating inter-protein EET from CP26 to LHCII. In the lifetime range of 3-15 ps, an excitation population growth is observed in CP43 while the excitation population decays in LHCII, indicating EET from LHCII to CP43. Within the same timescale, the dynamics involving CP26 are more complex, showing an excitation frequency-dependence. With 15,200-15,400 cm^-1^ excitation, the CP26 LDM shows growth, and the only corresponding decay is found in LHCII, suggesting EET from LHCII to CP26. In the 15,400-15,600 cm^-1^ region, the CP26 LDM shows instead a decay, with the only corresponding growth found in CP43, suggesting EET from CP26 to CP43. Overall, around 1-3 ps, energy flows from CP26 to LHCII. Around 3-15 ps, both CP26 (15,400-15,600 cm^-1^) and LHCII (15,200-15,600 cm^-1^) transfer energy to CP43 (Figure 2p).

In summary, 2DEV spectroscopy combined with the kinetic model constructed by the structure-based simulation show that the excitation of Chls *a* leads to initial EET from the core to the external antennae, while excitation of Chls *b* is followed by EET from the external antennae to the core, with additional inter-protein EET pathways between CP26 and LHCII. In the following, we provide a more detailed discussion of the kinetic model proposed in this work.

## Detailed analysis of the kinetic model

2DEV spectroscopy raises the prospect of the extraction of inter-protein EET dynamics experimentally. However, the large number of Chls in the PSII-SC still causes significant spectral congestion, obscuring the excitation population evolution of certain proteins. For example, the evolution of the peripheral antennae is not observed in the lower-frequency excitation range (14,700-15,200 cm^-1^) of the 2DEV spectra. Instead, the spectra are dominated by the evolution of CP43. This is because the excitation population in a protein needs to undergo enough evolution for the dynamics to be extracted experimentally. Figure 3 shows the simulated energy distribution at different time delays in the PSII-SC at two selected excitation frequencies: 14,800 cm^-1^, representing Chls *a* excitation (Figure 3a), and 15,400 cm^-1^, representing Chls *b* excitation (Figure 3b). The excitation distribution evolution clearly shows that the challenges of extracting inter-protein EET in the PSII-SC from the experiment originate not only from the significant spectral congestion but also from the intrinsic dynamics of the system. At 14,800 cm^-1^, simulations show that only the core antennae undergo obvious population evolution, while, being already partially excited, the population in the peripheral antennae barely changes. Therefore, it is expected that CP43 signatures on the 2DEV spectra show more evolution compared to the peripheral antennae in this excitation frequency range. Similarly, the simulated evolution of the excitation population shows that LHCII changes the most upon excitation at 15,400 cm^-1^, supporting the experimental observation that LHCII signatures show more dynamical evolution on the 2DEV spectra in this excitation frequency range and obscure the dynamics of the other protein subunits. These predictions made by the structure-based simulation completely match the observation from the 2DEV experiments, showing that the nature of the dynamic evolution can pose an additional constraint for experimental analyses, particularly in complex systems such as the PSII-SC.

**Fig. 3.**
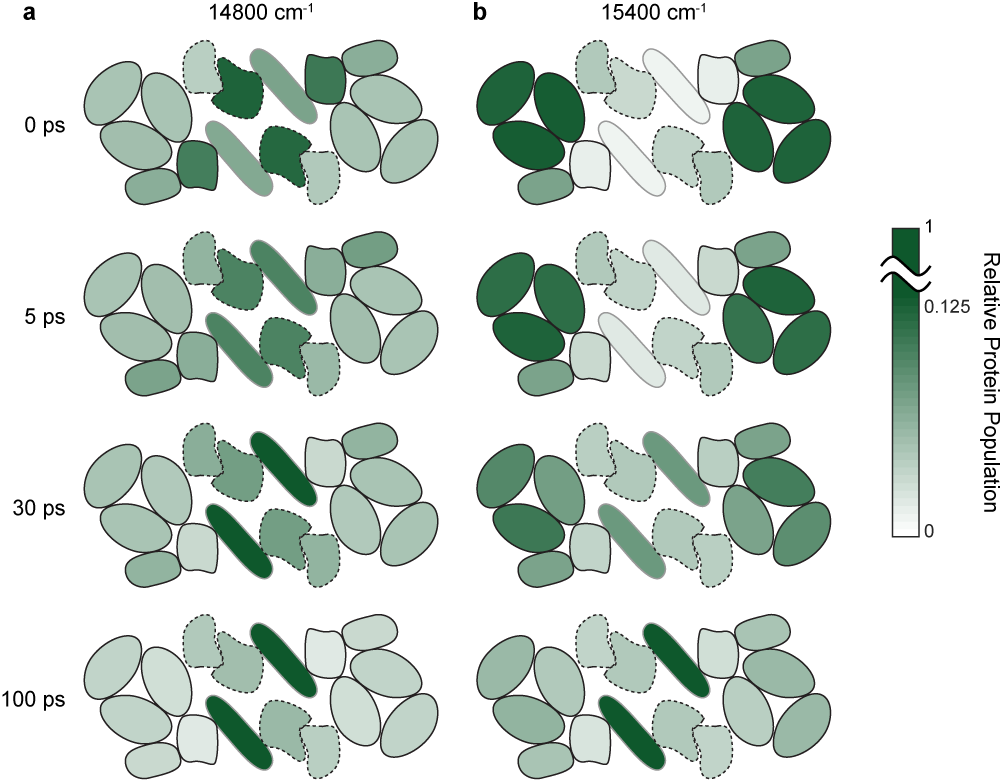
Simulated energy distributions at 77 K. Simulated excitation population in each protein (normalized to the total population) at 0, 5, 30, and 100 ps for **a**, 14,800 cm^-1^ and **b**, 15,400 cm^-1^. The arrangement of the protein subunits is labeled in Figure 1b. The protein subunits on the D1 side are represented by solid lines, and those on the D2 side are represented by dashed lines. The green scale indicates the relative excitation population in each protein (with the maximum in the color scale being 12.5% of the total population to provide visual enhancement for the difference).

Another important factor that needs to be addressed is that only the subunits with detectable IR signatures can be tracked experimentally via 2DEV spectroscopy. This limits the amount of information retrievable for CP47, which has been shown to have weaker IR features than CP43 on the 2DEV spectra of the PSII-CC ^39^. To understand the EET dynamics on the D2 side, at present, we can only rely on simulations that show great consistency with the experimental results for the dynamics on the D1 side. Figure 4a-j shows the LDM for the simulated population evolution of each protein in the PSII-SC at all excitation frequencies discussed (14,600-15,600 cm^-1^). The kinetic model shows that the EET directions of the proteins along the D2 branch, CP47 and CP29 (Figure 3d-e), are similar to the corresponding D1 proteins, CP43 and CP26 (Figure 4a-b, respectively. The excitation of Chls *a* leads to inter-protein EET from CP47 to CP29 while the excitation of Chls *b* shows EET from CP29 to CP47, consistent with the excitation frequency-dependent EET directions between the core and peripheral antennae observed for the D1 side. Both processes take place in 3-10 ps. The most significant difference is the 0.2-1 ps inter-protein EET between CP43 and LCHII/CP26 observed in the D1 proteins in the lower-frequency excitation range, which is not observed for the proteins on the D2 side.

**Fig. 4.**
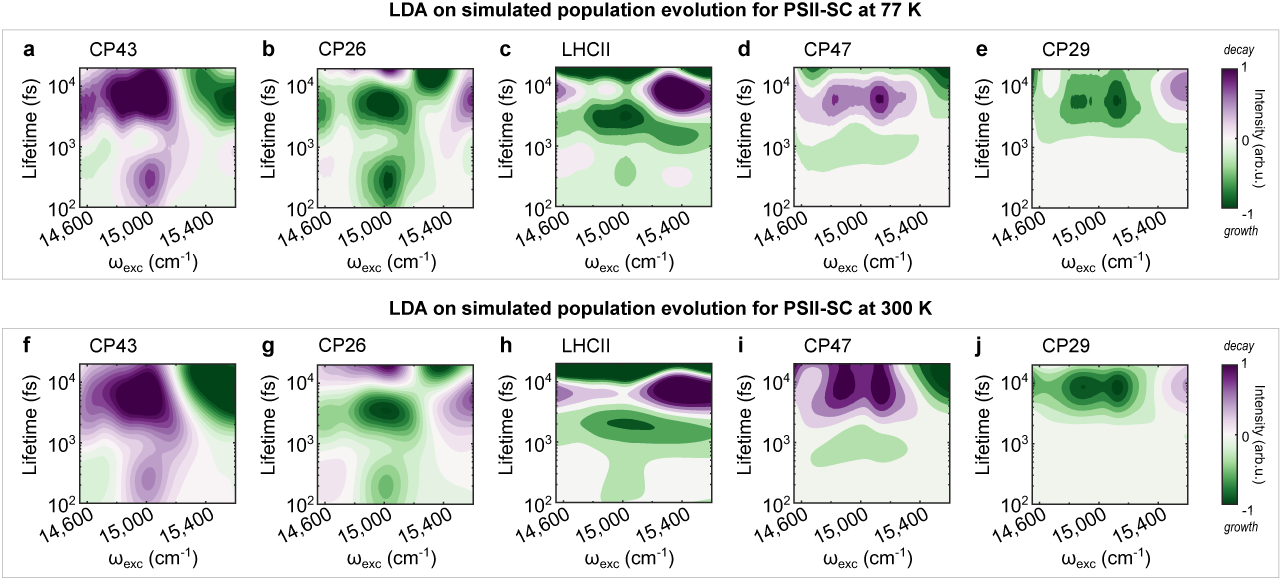
Excitation-dependent LDMs at 77 K and 300 K for the simulated population evolution of each protein in the PSII-SC. Simulated PSII-SC excitation-dependent LDM of **a**, CP43, **b**, CP26, **c**, LHCII, **d**, CP47 and **e**, CP29 at 300 K for all the excitation frequency ranges discussed. Simulated PSII-SC excitation-dependent LDM of **f**, CP43, **g**, CP26, **h**, LHCII, **i**, CP47 and **j**, CP29 at 300 K for all the excitation frequency ranges discussed. Positive amplitudes (purple) indicate decay; negative amplitudes (green) indicate growth.

Finally, the experiments were performed at 77 K to avoid photodamage. This raises a question about the applicability of the experimental data and the conclusion we can draw from them about natural photosynthesis. To understand the effect of temperature, simulations of the dynamics at 300 K were also performed. The model shows that the dynamics at 77 K (Figure 4a-e) and 300 K (Figure 4f-j) are almost identical within the first 20 ps. This confirms that the experiments performed at cryogenic temperatures can also provide insights relevant to physiological conditions.

In summary, combining 2DEV spectroscopy and theoretical simulation enables the understanding of the EET dynamics in the PSII-SC, which allows us to conclude that initial EET involves transfer from the core to the external antennae upon excitation of Chls *a* and from the external antenna to the core upon excitation of Chls *b*. These observations are a good representation of the important EET pathways present in the native environment and have a significant impact on the functions of PSII-SC. In particular, the bidirectional EET has impacts on the charge separation efficiency at low light levels and photo-protection in excess light. We discuss this point in more detail in the following section.

## Efficiency and photoprotection

In the previous sections, we showed that, in the lower-frequency excitation range, the energy quickly leaves the core to explore the peripheral antennae, faster than it is transferred to the RC. The ability of the former process to compete with the latter is crucial for photoprotection. Indeed, it has been shown that photoprotection mostly takes place within the peripheral antennae via interactions with carotenoids ^2,25,47,48^. On the other hand, for the higher-frequency excitation range, in which mostly Chls *b* (found only in the peripheral antennae) are excited, the net EET direction is the opposite. The absorbed energy, mostly distributed in the external antennae upon excitation, has already a higher chance to explore the quenching sites due to their proximity to the initially excited pigments, which leads to effective protection. Under low light conditions, where the quenchers are inactive, the EET pathways quickly guide the unquenched energy from the external antennae toward the core to reach the RC.

One key factor that allows the bidirectionality of the energy flow, which facilitates the switch between efficient and photoprotective mode, is the timescale of different competing EET pathways. In the lower-frequency excitation range, energy is observed to transfer from CP43 to CP26 and LHCII on a sub-ps timescale. This pronounced connection between the core and the peripheral antennae on the D1 side is a result of the short center-to-center distances between certain pairs of Chls of CP26/LHCII and CP43, e.g., C601-C513 (CP26-CP43: 12.6 Å), C614-C503 (CP26-CP43: 16.02 Å), and C611-C506 (LHCII-CP43: 17.05 Å) ^13^. This kind of design allows the EET pathways that guide energy out of the core to compete with EET from CP43 to the RC, which has been shown to happen on a timescale of 10s of ps ^20,26,30,39,46^. Such ultrafast EET between different subunits was not observed in the simulated population evolution of the subunits along the D2 side (Figure 4d-e,i-j). However, within 3-10 ps, EET from the core to the peripheral antennae is observed along both sides. This still allows EET from CP47 to CP29 to compete with EET from CP47 to the RC as the latter has been shown to be slower compared to EET from CP43 to the RC ^30^. On the other hand, in the higher-frequency excitation range where excitation is further away from the RC, inter-protein EET to the core antennae still occurs in 3-15 ps. This is a similar timescale to the EET from the core antenna to the RC, which guarantees that energy reaches the RC before it is dissipated.

To illustrate the significance of the actual timescales of energy flow in the PSII-SC, we make use of a conceptual coarse-grained model (Figure 5a). By employing the timescales observed in our experiments in the coarse-grained model, we demonstrate how the balance between efficient charge separation and photoprotection rely on the EET timescales between peripheral and core antennae. Figure 5b,c shows that, under natural operation conditions of the PSII-SC, the population of the peripheral antennae (blue) grows faster than that of the RC (black), allowing energy to visit the quenching sites before entering the RC. A five-fold slower EET from the core antennae to the peripheral antennae (Figure 5d,e) would greatly reduce the probability of visiting the quenching sites because the transfer to the RC (black) would become dominant as the population of the peripheral antennae (blue) is suppressed. This would limit the ability of the PSII-SC to quench excessive excitation under high-light conditions. On the other hand, a five-fold slower transfer from the peripheral to the core antennae (Figure 5f,g) would decrease the photosynthetic efficiency in low-light conditions as energy cannot fully reach the RC (black) before undergoing dissipative pathways. Such a balance shows that the structural arrangement of the peripheral and the core antennae allows the PSII-SC to work in a regime where EET occurs on timescales optimized for both efficiency and photoprotection. Noticeably, the timescales found in this work are designed to be balanced with the EET timescale from the core antennae to the RC, which is limited by the large distance required for protecting the separated charges ^49^. Furthermore, the effect of transfer timescales from the peripheral to the core antennae shows that there is an upper limit for the antenna size (larger than the C_2_S_2_-type PSII-SC), as proposed by Croce and coworkers ^50^. An increased antenna size leads to an increased overall absorption cross-section, which is necessary under low-light conditions. However, it also leads to slower EET to the PSII-CC, imposing an upper limit on the antenna size for optimal photosynthetic efficiency.

**Fig. 5.**
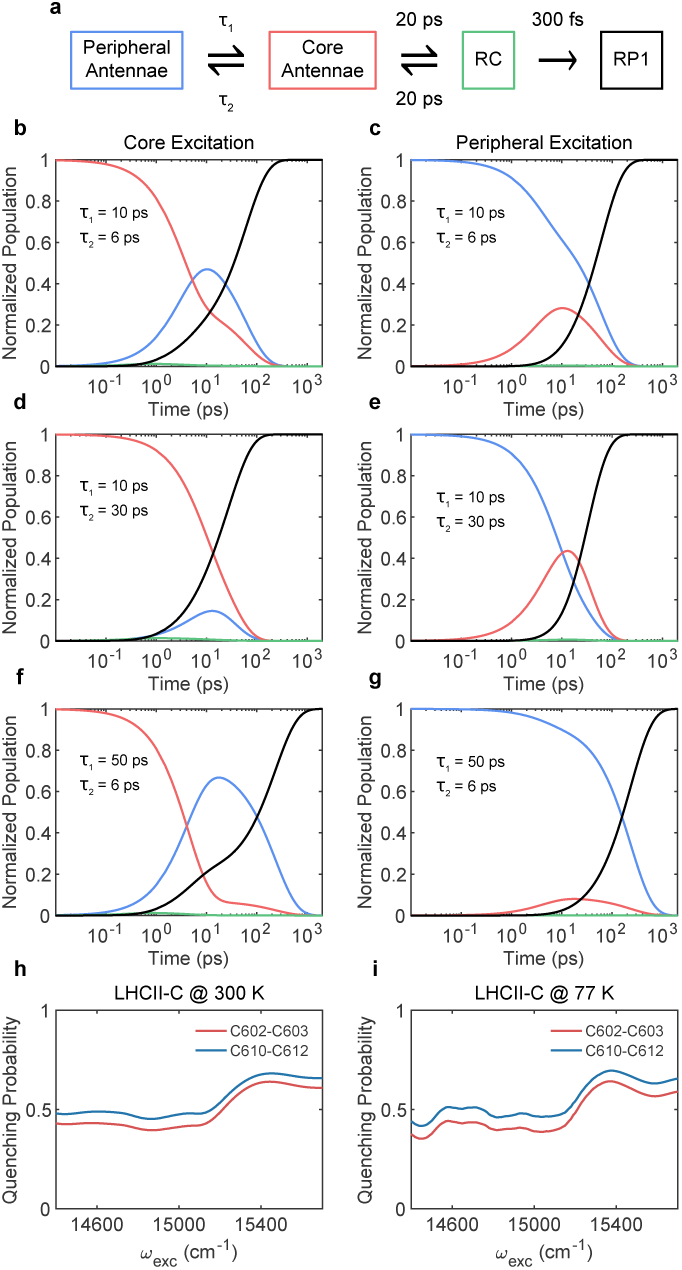
A coarse-grained kinetic model for the bidirectional EET between core and peripheral antennae. **a**, A coarse-grained kinetic model with the timescales approximated from the simulation and literature ^20,26,30,39,46^). Population evolution upon the excitation of core/peripheral antennae, respectively, when the timescales are **b,c**, extracted from the 2DEV measurement **d,e**, five-fold slower for the core to peripheral antennae transfer and **f,g**, five-fold slower for the peripheral to core antennae transfer. The color scheme of **b-g** is consistent with the compartments in **a**. Simulated excitation-dependent quenching probability when active quenchers are placed in the LHCII-C monomer (see Figure 1) for **h**, 300 K and **i**, 77 K. The quenchers are placed in proximity to C602-C603 (red) and C610-C612 clusters (blue). A detailed description can be found in Extended Data Figure E.3 and in Methods.

Overall, the kinetic network in the PSII-SC is designed so that, regardless of where excitation is in the PSII-SC, excitation energy has a high chance of visiting the quenching sites. There, it can either be quenched in high-light conditions or continue to be transferred to the RC well before non-radiative processes occur in low-light conditions. Figure 5h, i (and Extended Data Figure E.3) shows that, by placing the quencher in the peripheral antenna (e.g. LHCII), the protection ability of the quencher does not depend on the excitation axis, and therefore does not depend on the excitation location. This shows that non-photochemical quenching, which uses a feedback loop to activate/deactivate quenching depending on light intensity ^2^, can only work in combination with the bidirectional EET pathways with balanced timescales. In other words, a fine kinetic balance is necessary to allow effective photoprotection under high-light conditions (quenching activated) while guaranteeing photosynthetic efficiency under low-light conditions (quenching inactive). How the balance (Figure 5b, c) is achieved depends on the detailed microscopic rates of energy flow between the subunits of the PSII-SC that are obtained by the comparison of experimental and simulation data described above.

## Concluding remarks

It has been known for several decades that the PSII-SC has a rather flat energy landscape ^3–7^, in contrast to the energy funnel that other photosynthetic systems exhibit ^1^. While it is speculated that the flat energy landscape (or shallow funnel) design is related to photoprotection ^4,7^, the exact working mechanism cannot be easily inferred without a deeper understanding of the EET dynamics in the PSII-SC.

In this work, we reveal the connection between the EET network and the functions of the C_2_S_2_-type PSII-SC. We show that energy can flow out of the PSII-CC and transfer from the peripheral antenna system back into the core, increasing the probability of visiting the quenching sites before entering the RC. The timescales for the net EET between peripheral and core antennae, controlled by the microscopic transfer rates, are finely balanced to facilitate the bidirectionality of energy flow in the PSII-SC. It is reasonable to argue that such an optimized kinetic network is made possible by the rather flat energy landscape within the multi-component structure of the PSII-SC. Ultimately, the strategy to have bidirectional energy transfer pathways enables switching between efficient energy conversion and effective photoprotection, responding to the fluctuating sunlight intensity.

Understanding the functional mechanism of the PSII-SC can allow us to improve the bio-inspired design of solar energy devices, enabling fine control of the solar energy conversion processes. Additionally, the optimization of the response of crop plants to fluctuating light levels has emerged as a major step in the improvement of crop yield ^10^. As the location and timescales associated with non-photochemical quenching are elucidated, the detailed understanding of energy flow pathways and timescales, as well as the response to differing solar wavelengths, will aid in formulating the strategies to continue to enhance the yields of food crops, as is necessary over the next 20-30 years.

## Supplementary information

Supplementary Information is available for this paper.

## Acknowledgments

This research was supported by the US Department of Energy, Office of Science, Basic Energy Sciences, Chemical Sciences, Geosciences, and Biosciences Division. We thank Prof. Doran I. G. B(ennett) Raccah for sharing the code for simulation and providing useful suggestions.

## Declarations

### Ethics approval

Not applicable

### Consent to participate

Not applicable

### Consent for publication

Not applicable

### Availability of data and materials

The data presented in this work are available from the corresponding author upon request.

### Code availability

The codes used in this work are available from the corresponding author upon request.

### Authors’ contributions

E.A.A. and G.R.F. conceived the study design. C.L., S.-J.Y. and K.O. performed the 2DEV experiments. C.L. performed LDA. S.-J.Y. performed the theoretical simulations. M.I. prepared the sample. C.L., S.-J.Y. and G.R.F. wrote the paper. All author commented on the manuscript.

## Methods

### Sample Preparation

All procedures for sample preparation were performed in the dark to minimize exposure to light as much as possible. We prepared PSII-enriched membranes from spinach leaves according to the previous literature ^1,2^ with some modifications as described previously ^3,4^. For preparing the C_2_S_2_-type PSII-SC, the PSII-enriched membranes (0.5 mg Chl/mL) were solubilized with 1.0% (w/v) *α*-DDM (n-dodecyl-*α*-D-maltopyranoside, Anatrace) in a buffer containing 25 mM MES-NaOH (pH 6.0) for 30 min on ice. The solubilized membranes were then centrifuged at 21,000 × g for 5 min at 4*^◦^*C. The supernatants were loaded onto sucrose gradients (each concentration overlaid with the denser one: 2.1 mL of 0.1, 0.4, 0.7, 1.0, and 1.3 M sucrose and 0.5 mL of 1.8 M sucrose at the bottom) in ultracentrifuge tubes (14 × 89 mm, Beckman Coulter). The sucrose gradients contained 0.03% (w/v) *α*-DDM buffered as above. Centrifugation was performed at 154,300 × g for 24 h at 4*^◦^*C (SW 41 Ti swinging-bucket rotor, Beckman Coulter). The separated bands were collected dropwise from the bottom of the tube. The collected fraction containing the C_2_S_2_-type PSII-SC was concentrated using a centrifugal filter (100K MWCO). The concentrated sample was diluted with a buffer containing 25 mM MES-NaOH (pH 6.0), 10 mM NaCl, 3 mM CaCl_2_, 400 mM sucrose, and 0.03% *α*-DDM prepared with D_2_O. The concentrated C_2_S_2_-type PSII-SC was flash-frozen and stored at -80*^◦^*C until 2DEV measurements. The preparation of isolated LHCII follows the procedure in the work of Arsenault et al. ^5^ with the only exception being the final resuspension step, which is done with the same buffer used in the preparation of the C_2_S_2_-type PSII-SC.

### 2DEV Measurements

The details of 2DEV set up can be found elsewhere ^3,4,6^. The reported PSII-SC 2DEV data were collected in two separate measurements, both at 77 K (Extended Data Figure E.1). For the first measurement, the excitation pulses were centered at 665 nm with FWHM of 70 nm, and the sample was diluted with glycerol to have an optical density of *∼*0.6 at 675 nm. For the second measurement, the excitation pulses were centered at 630 nm with FWHM of 55 nm, and the sample was diluted to have an optical density of *∼*1.0 at 650 nm. For isolated LHCII, the 2DEV measurement was performed at 77 K with excitation pulses centered at 655 nm with FWHM of 55 nm. The sample was diluted with glycerol to have an optical density of *∼*0.8 at 675 nm. For all measurements, the optical path length was *∼*200 *µ*m and the excitation pulses were compressed to 15*∼*20 fs. Both pulses were focused to a spot size of *∼*200 *µ*m with a combined energy *∼*80 nJ. The detection pulse was centered at 5,900 nm and spanned over 5,500*∼*6,300 nm. The instrument response is estimated to be *∼*90 fs from cross-correlation between the visible and mid-IR pulses. The time delays between the second visible pump and the IR probe are the same for all measurements: from 0 to 1.04 ps with steps of 20 fs; from 1.14 to 20 ps with steps of 100 fs; from 25 to 100 ps with steps of 5 ps. The repetition rate of the source laser is set to 1 kHz.

### Lifetime Density Analysis (LDA)

LDA approximates the data evolution as a sum of hundreds of exponentials to retrieve the corresponding amplitudes (*x_j_*(*τ_j_, λ*)), producing lifetime density maps (LDM) ^7^:

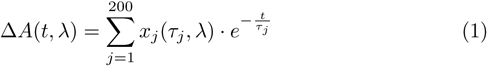

Despite LDA resulting in high uncertainty in the time constants, the ability to handle hundreds of exponential trends with little initial assumptions is an important advantage over the widely used global analysis ^8^. The data analysis was performed using pyLDM ^9^. Regularization of the minimization process is always applied to obtain reliable LDMs ^9^. A low regularization hyperparameter *α*, which corresponds to higher LDM amplitudes and narrower lifetime distributions, is initially adopted. However, low values of *α* have a higher chance to return artifacts. This is true independently of the noise level, as it is observed also for the noise-free simulated population evolution. Extended Data Figure E.2a shows an example of LDM obtained for a low *α*, showing that the artifact appears as a satellite peak with opposite amplitude compared to the main lifetimes. To identify the artifacts, exponential fits of the LDM obtained for the regularization parameter *α* = 0.1 are performed (see Supplementary Information, Extended Data Figure E.2 and Extended Data Table 3). Based on the fitting, we select the optimal hyper-parameter *α* that returns artifact-free LDMs. In particular, we find that *α* = 3 provides the best results for the LDA of all experimental data and simulated population evolution (Extended Data Figure E.2b). Only for the simulated LHCII excitation population an *α* = 1 was adopted. We note that a higher hyper-parameter *α* leads to wider lifetime distributions (Extended Data Figure E.2a-c). The broadening of the lifetime distributions, therefore, is merely a consequence of the analysis ^9^.

### EET Dynamics Simulation

The simulations were performed based on a modified version of the structure-based model proposed in the work of Bennett et al., where detailed simulation procedures can be found ^10^. The differences between the model used in this work and the work of Bennett et al. are listed and discussed here. The parameters used for the simulations can be found in the Extend Data Table 1 and 2.

1. The inter-protein pigment couplings were calculated based on the cryo-EM structure of the C_2_S_2_-type PSII supercomplex extracted from spinach (PDB: 3JCU) ^11^, and the TrEsp method was used instead of applying point dipole approximation ^12^. The atomic transition charges of Chls *a*, Chls *b* and pheophytins a were obtained from literature ^12–14^, and scaled to match the transition dipole moments listed in Extended Data Table 1.
2. CP29 Hamiltonian, originally represented by the LHCII monomer Hamiltonian in Bennett’s model, was described by a new Hamiltonian proposed by Mascoli et al. ^15^ The new CP29 Hamiltonian was constructed based on isolated CP29, in which C616 is absent due to purification. Therefore, the C616 is not included in the CP29 Hamiltonian. Additionally, the 13-state Hamiltonian contains C614, which is absent in the 3JCU structure and is, therefore, deleted from the Hamiltonian. Currently, there is no semi-empirical Hamiltonian for CP26. Due to the spectral similarity between CP26 and CP29 ^16^, the CP26 Hamiltonian is represented by the CP29 Hamiltonian with C614 included, which is present in CP26 in the 3JCU structure.
3. The line-broadening functions were calculated for 77 K to match the experimental conditions. Unlike the calculation for 300 K (both in this work and Bennett’s model), the lineshape functions for the core components (RC, CP43 and CP47) do not converge in the time domain without a dephasing term, which was originally included in the work of Renger and coworkers ^17,18^ but omitted in Bennett’s model. At 300 K, the effect of the dephasing term is negligible due to stronger electron-phonon interaction, and can therefore be omitted. For the calculations at 77 K, the term is required to ensure the convergence of lineshape functions. In our simulations, the dephasing term was included for the core components and the dephasing time was taken as 1 ps. Different values were tested and the effect is negligible compared to inhomogeneous broadening.

The population of each state at each time point was then calculated based on the hybrid rate matrix (combining generalized-Förster and modified-Redfield rates, see ref. ^10^) with the following equation:

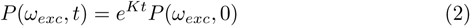

where *P* (*ω_exc_, t*) is the excitation-dependent population, *K* is the hybrid rate matrix and *P* (*ω_exc_,* 0) is the excitation frequency-dependent initial population calculated based on the integrated absorption of individual excitonic states within a 10 cm^-1^ range (centered at each defined excitation frequency). The population evolution of each protein was calculated by converting the exciton population to the Chl population and summing over all the Chls within each protein. Simulation of single excitation (Figure 1c-h) was performed by assigning excitation to one single Chl as the initial population, C509 for CP43 and C611 for CP47, both of which are at the center of each protein. We note that, in principle, the charge separation dynamics can be included in the rate matrix by incorporating the timescales obtained from the fitting of experimental data. However, such fitting has been demonstrated to be problematic as different models can provide equally good fits ^10^. Therefore, instead of including empirical charge separation lifetimes in our model, it is assumed that charge separation occurs infinitely faster than the EET out of RC components. Such an approximation is not only consistent with the transfer-to-trap limited model ^18–20^ reported in the literature but it has also been applied to another model ^21^ which was able to reproduce experimental results with excellent agreement. While we do not expect the approximation to change the overall dynamics of energy transfer, especially at early waiting times, future improvements can be made by properly incorporating descriptions of charge transfer dynamics into the model.

We note that Bennett et al. also proposed the “domain model”, in which it is assumed that intra-domain EET is fast enough to allow thermal equilibrium within each domain before inter-domain EET occurs. They showed that the dynamics predicted by the hybrid model and the domain model share great similarities. However, in our calculations, the dynamics differ dramatically when thermal equilibrium is assumed at the cryogenic condition (77 K). In contrast, the room temperature (300 K) simulations, which most likely allow faster equilibrium, generate similar results with both models, as described in the work of Bennett et al. For consistency, the population evolution of both conditions was calculated based on the hybrid rate matrix instead of the domain-to-domain transfer rate matrix.

The simulation of quenching probability (Extended Data Figure E.3) with activated quenchers was performed by connecting additional sinks (where reverse transfer is prohibited) to the Chls that are suspected of being responsible for EET to carotenoids, C602-C603 and C610-C612 ^22,23^. The rate for EET from these Chls to carotenoid is set to be (200 fs)^-1^, which is similar to the values reported in literature ^24,25^. The quenching probability is then defined as the ratio between the population in the sinks and the RC at the long time limit. In each simulation, two quenchers were each placed in the same protein subunit of the two PSII-SC monomers. All simulations were performed for both 77 K and 300 K.

---

**Extended Data Figure 1.**
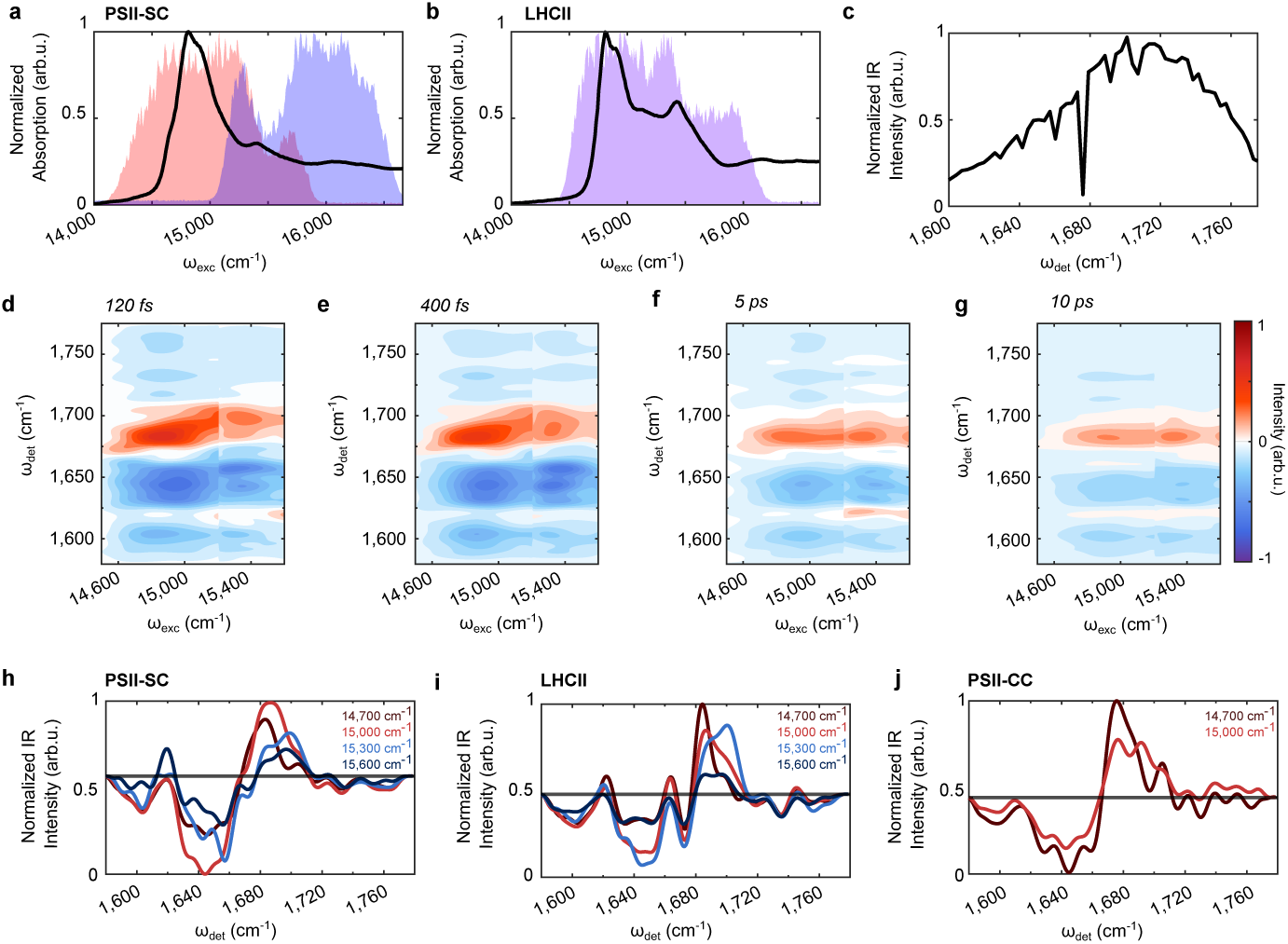
Experimental spectra. Normalized absorption spectrum and visible excitation pump spectra for the 2DEV measurements on the **a**, PSII-SC and **b**, isolated LHCII trimer. **c**, Infrared probe spectrum used for all 2DEV measurements. 2DEV maps at different time delays for the PSII-SC: **d**, 120 fs, **e**, 400 fs, **f**, 5 ps and **g**, 10 ps. The excitation at 15,200 cm^-1^ marks the separation between the two PSII-SC measurements (see panel (a) and Methods). 2DEV spectral slices at 120 fs normalized between 1 and -1 for excitation frequencies of 14,700 cm^-1^ (dark red), 15,000 cm^-1^ (red), 15,300 cm^-1^ (blue) and 15,600 (dark blue) cm^-1^ for **h**, PSII-SC, **i**, isolated LHCII and **j**, PSII-CC. The 2DEV slices of the PSII-CC were reproduced with permission from Yang et al. Proc. Natl. Acad. Sci. U.S.A. 119, e2208033119 (2022). Copyright (2022) National Academy of Sciences.

**Extended Data Figure 2.**
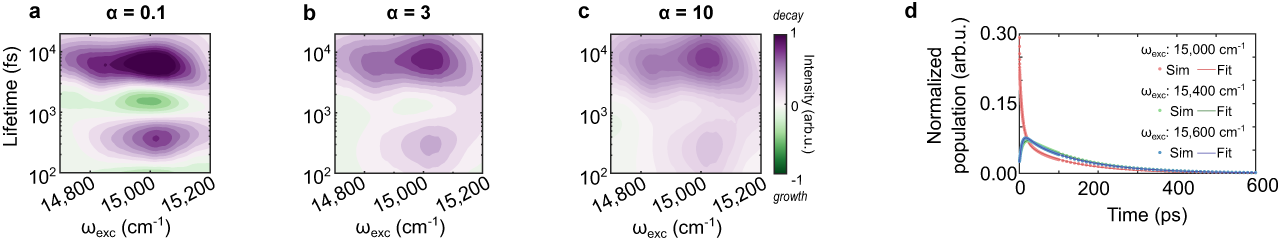
Example of cross-checking of LDA analysis results. LDM for the simulated population evolution of CP43 in the PSII-SC for hyper-parameter *α* of **a**, 0.1, **b**, 3 and **c**, 10. At increasing *α* the negative amplitude around 1 ps disappears. The corresponding growth is not observed via exponential fitting of the simulated population evolution of CP43 at different excitation frequencies (see Extended Data Table 3 and Supplementary Information). **d**, Exponential fit of the simulated population evolution of CP43 for three different excitation frequencies (fit parameters are reported in Extended Data Table 3)

**Extended Data Figure 3.**
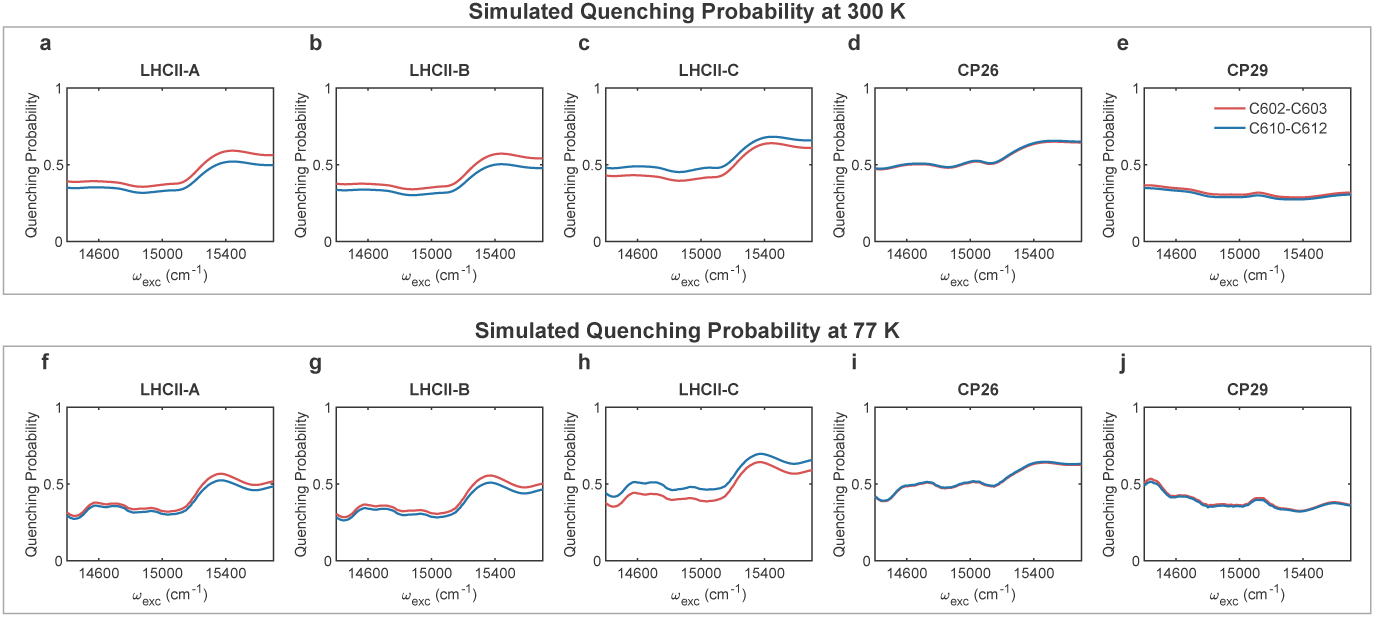
Simulated quenching probability in the peripheral antennae of the PSII-SC at 300 K and 77 K. Probability that the excitation energy is quenched before reaching the RC, with quenching sites (the Chls near carotenoids) being in (a, f) LHCII-A (G/g), (b, g) LHCII-B (N/n), (c, h) LHCII-C (Y/y), (d, i) CP26 and (e, j) CP29 at 300 K (a-e) and 77 K (f-j). The EET rate from Chls to carotenoid is universally set to (200 fs)^-1^ and the exact quenching sites (red: C602-C603; blue: C610-C612) are selected based on literature. ^22,23^ Detailed description can be found in Methods.

**Extended Data Table 1.** EET simulation parameters.

|  | RC<sup>1</sup> |  | CP43<sup>2</sup>/CP47<sup>3</sup> | CP26/CP29<sup>4</sup> |  | LHCII<sup>5</sup> |  |
| --- | --- | --- | --- | --- | --- | --- | --- |
| Pigment | Chl *a* | Pheo | Chl *a* | Chl *a* | Chl *b* | Chl *a* | Chl *b* |
| TDM<sup>6</sup> | 4.4 | 3.5 | 4.4 | 3.74 | 3.18 | 4 | 3.4 |
| $\sigma_{\text{inhom}}$<sup>7</sup> | 200 | 120 | 180 | 90 | 108 | 80 | 96 |
| SD type<sup>8</sup> | a | b | c | d | e | f | g |

<sup>1</sup>Raszewski et al. 17,18 (adapted by Bennett et al. 10)
<sup>2</sup>Müh et al. 26 (adapted by Bennett et al. 10)
<sup>3</sup>Raszewski et al. 18 (adapted by Bennett et al. 10)
<sup>4</sup>Mascoli et al. 15
<sup>5</sup>Novoderezhkin et al. 27 (adapted by Bennett et al. 10)
<sup>6</sup>Transition dipole moment magnitude [unit: Debye]
<sup>7</sup>Inhomogeneous broadening width [unit: $\text{cm}^{-1}$ ]
<sup>8</sup>Spectral density type in Extended Data Table 2

**Extended Data Table 2.** Spectral density parameters (detailed description can be found in Supplementary Information).

|  | a | b | c | d | e | f | g |
| --- | --- | --- | --- | --- | --- | --- | --- |
| $S_0$ | 0.65 | 0.65 | 0.5 | | | | |
| $s_1$ | 0.8 | 0.8 | 0.8 | | | | |
| $s_2$ | 0.5 | 0.5 | 0.5 | | | | |
| $\omega_1$ [cm<sup>-1</sup>] | 0.532 | 0.532 | 0.532 | | | | |
| $\omega_2$ [cm<sup>-1</sup>] | 1.94 | 1.94 | 1.94 | | | | |
| $\lambda_0$ [cm<sup>-1</sup>] | | | | 40 | 48 | 37 | 48 |
| $\gamma_0$ [cm<sup>-1</sup>] | | | | 40 | 40 | 30 | 30 |
| UD BO<sup>1</sup> | No | No | No | Yes | Yes | Yes | Yes |

<sup>1</sup>This row indicates whether under-damped Brownian oscillators were included in the spectral density.

**Extended Data Table 3.** Exponential fit of simulated population evolution of the PSII-SC subunits. A detailed description can be found in Supplementary Information.

| $\omega_{exc}$ [cm<sup>-1</sup>] | CP43 | | | CP26 | | |
| --- | --- | --- | --- | --- | --- | --- |
|  | 15000 | 15400 | 15600 | 15000 | 15400 | 15600 |
| $A_1$ | 0.07 | -0.06 | -0.11 | -0.06 | 0.01 | 0.05 |
| $\tau_1$ (ps) | *0.36* | *8.02* | *7.74* | *0.32* | *1.20* | *7.57* |
| $A_2$ | 0.15 | 0.09 | 0.05 | -0.06 | -0.01 | 0.05 |
| $\tau_2$ (ps) | *8.24* | *149.20* | *15.23* | *5.86* | *4.93* | *57.24* |
| $A_3$ | 0.07 | | 0.08 | 0.10 | -0.04 | 0.13 |
| $\tau_3$ (ps) | *123.20* | | *147.80* | *33.18* | *20.60* | *159.30* |
| $A_4$ | | | | 0.10 | 0.17 | |
| $\tau_4$ (ps) | | | | *163.00* | *145.90* | |
| $R^2$ | 0.99974 | 0.99984 | 0.99974 | 1.00000 | 0.99997 | 0.99999 |

| $\omega_{exc}$ [cm<sup>-1</sup>] | CP47 | | | CP29 | | |
| --- | --- | --- | --- | --- | --- | --- |
|  | 14900 | 15300 | 15600 | 14900 | 15300 | 15600 |
| $A_1$ | 0.04 | 0.04 | -0.08 | -0.07 | -0.04 | 0.08 |
| $\tau_1$ (ps) | *6.50* | *5.71* | *11.56* | *5.90* | *6.01* | *11.71* |
| $A_2$ | 0.17 | -0.12 | -0.09 | 0.14 | -0.18 | -0.12 |
| $\tau_2$ (ps) | *183.40* | *73.66* | *65.21* | *170.90* | *180.20* | *108.90* |
| $A_3$ | | 0.21 | 0.20 | | 0.26 | 0.23 |
| $\tau_3$ (ps) | | *175.10* | *175.40* | | *156.00* | *161.30* |
| $R^2$ | 0.99932 | 0.99996 | 0.99998 | 0.99981 | 0.99988 | 0.99997 |

| $\omega_{exc}$ [cm<sup>-1</sup>] | LHCII | | | RC | | |
| --- | --- | --- | --- | --- | --- | --- |
|  | 15000 | 15400 | 15500 | 14800 | 15000 | 15400 |
| $A_1$ | -0.01 | -0.01 | 0.02 | -0.06 | -0.10 | 0.04 |
| $\tau_1$ (ps) | *0.51* | *1.31* | *10.23* | *10.52* | *34.50* | *10.08* |
| $A_2$ | -0.04 | 0.06 | 0.24 | -0.80 | -0.77 | -1.025 |
| $\tau_2$ (ps) | *4.48* | *9.15* | *102.20* | *159.60* | *169.70* | *166.00* |
| $A_3$ | -0.06 | 0.19 | 0.30 | 1.00 | 1.00 | 1.00 |
| $\tau_3$ (ps) | *28.57* | *70.51* | *172.10* | *3.76E5* | *3.39E5* | *3.13E5* |
| $A_4$ | 0.32 | 0.45 | | | | |
| $\tau_4$ (ps) | *160.60* | *158.10* | | | | |
| $R^2$ | 1.00000 | 1.00000 | 0.99999 | 0.99999 | 0.999980 | 1.00000 |

