## Supplementary Information for "Bidirectional Energy Flow in the Photosystem II Supercomplex"

### 1 Spectral Density Definitions

The spectral density parameters are listed in the Extended Data Table 2. Two different spectral densities were applied for the simulation to ensure consistency with the literature Hamiltonian of each protein subunit, as described by Bennett et al.[1]

For the PSII-CC components, the spectral density is defined[1,2] as

$$\chi''(\omega) = (\pi\hbar)\frac{S_0}{s_1 + s_2}\sum_{i=1,2}\frac{s_i\omega^5}{7!2\omega_i^4}e^{-\sqrt{\frac{\omega}{\omega_i}}} \tag{1}$$

where $S_0$, $s_1$, $s_2$, $\omega_1$, and $\omega_2$ are the parameters listed in the Extended Data Table 2.

For the minor antennae and LHCII, the spectral density is defined[1,3] as

$$\chi''(\omega) = 2\lambda_0\frac{\omega\Gamma_0}{\omega^2 + \Gamma_0^2} \tag{2}$$

where $\lambda_0$ and $\Gamma_0$ are the parameters listed in the Extended Data Table 2. Additionally, vibronic coupling with individual modes are also included for the peripheral antennae, which contributes to the spectral density as[1,3]

$$\chi''_{vib}(\omega) = \sum_{j=1}^{N_{vib}} 2S_j\omega_j^3\frac{\omega\Gamma_{vib}}{(\omega_j^2 - \omega^2)^2 + \omega^2\Gamma_{vib}^2} \tag{3}$$

where $S_j$, $\omega_j$, and $\Gamma_{vib}$ are the parameters for each vibration modes. The values of these parameters can be found in ref[1] and ref[4].

#### 2 Fitting of Simulated Excitation Population Evolution of the PSII-SC Subunits

The simulated excitation population evolution of PSII-SC subunits (see Methods) was subjected to exponential fitting with the following equation

$$f(t) = \sum_i A_i e^{\frac{t}{\tau_i}} \quad (4)$$

where $f(t)$ is the excitation population evolution (traces) of individual PSII-SC subunits, and $A_i$ and $\tau_i$ are the variables, reported in Extended Data Table 3 (see also Extended Data Figure 2). The number of components lies between 2-4, decided based on the LDA results as well as the fitting quality.

Fitting is necessary for confirming the validity of LDA, as artifacts rise for too low values of the regularization parameter $\alpha$ (see Methods Section 6.3 and Extended Data Figure 4), which influences the interpretation of both experimental and simulation results. The fittings were performed on the traces of selected excitation frequencies for each PSII-SC subunit. The excitation frequencies and initial parameters for the fittings are selected based on the LDMs obtained for $alpha = 0.1$. It is important to note that, in some cases, the traces cannot be described as the sum of a few exponential components. For example, the fitting results for CP47 traces display visible difference from the actual population evolution. Additionally, the fitting of the RC trace at 15,400 $cm^{-1}$ shows a decaying component around 10 ps. However, the population evolution of the RC should be monotonically increasing as it is assumed that charge separation occurs as soon as energy reaches the RC (instant trapping). These issues with fitting of noiseless simulated population evolution likely arise from the nature of non-exponential dynamics within the system. Since the system contains a large number of EET pathways, the complicated network does not always guarantee an exponential dynamics. Therefore, the lifetimes obtained from exponential fitting and LDA should be treated as a reflection of characteristic timescales, instead of the actual lifetimes of individual EET pathways.
